# A genetically buffered helicase network promotes tolerance of G-quadruplex stabilization in *Saccharomyces cerevisiae*

**DOI:** 10.64898/2026.07.07.737069

**Authors:** Spencer J. Gray, Matthew L. Bochman

## Abstract

G-quadruplexes (G4s) are non-canonical DNA secondary structures that can impede DNA replication and transcription and provoke genome instability, and DNA helicases of the PIF1 and RecQ families have long been regarded as the principal enzymes that resolve them. To directly test the relative contributions of these families, we measured the growth of *Saccharomyces cerevisiae* helicase mutants in the presence of the G4-stabilizing ligand pyridostatin (PDS). Unexpectedly, no single PIF1- or RecQ-family mutant was sensitized to PDS relative to wild type. Sensitivity emerged only in double mutants, and it did so for combinations both within a single family and across the two families. This pattern indicates that G4 tolerance is buffered by the combined, partially interchangeable, activity of multiple helicases rather than by any one family. To ask whether this redundancy extends beyond the canonical players, we tested two additional helicases whose human orthologs are implicated in G4 metabolism: Chl1 (DDX11/ChlR1) and Srs2 (RTEL1). Loss of Chl1 alone did not sensitize cells, and *chl1Δ* combined with PIF1- or RecQ-family mutations recapitulated the redundancy pattern – with one informative exception: *chl1Δ hrq1Δ* remained PDS-tolerant, placing Chl1 and Hrq1 in a shared genetic route. In contrast, *srs2Δ* was the sole single mutant sensitized to PDS, defining a non-redundant requirement that no other helicase compensates. We integrate these results into a two-layer model in which a redundant helicase pool resolves G4-associated genomic stress, while a non-redundant Srs2 function manages its recombinogenic consequences. Our findings reframe G4 maintenance from a family-specific activity into a distributed, buffered network.

**GRAPHICAL ABSTRACT:** 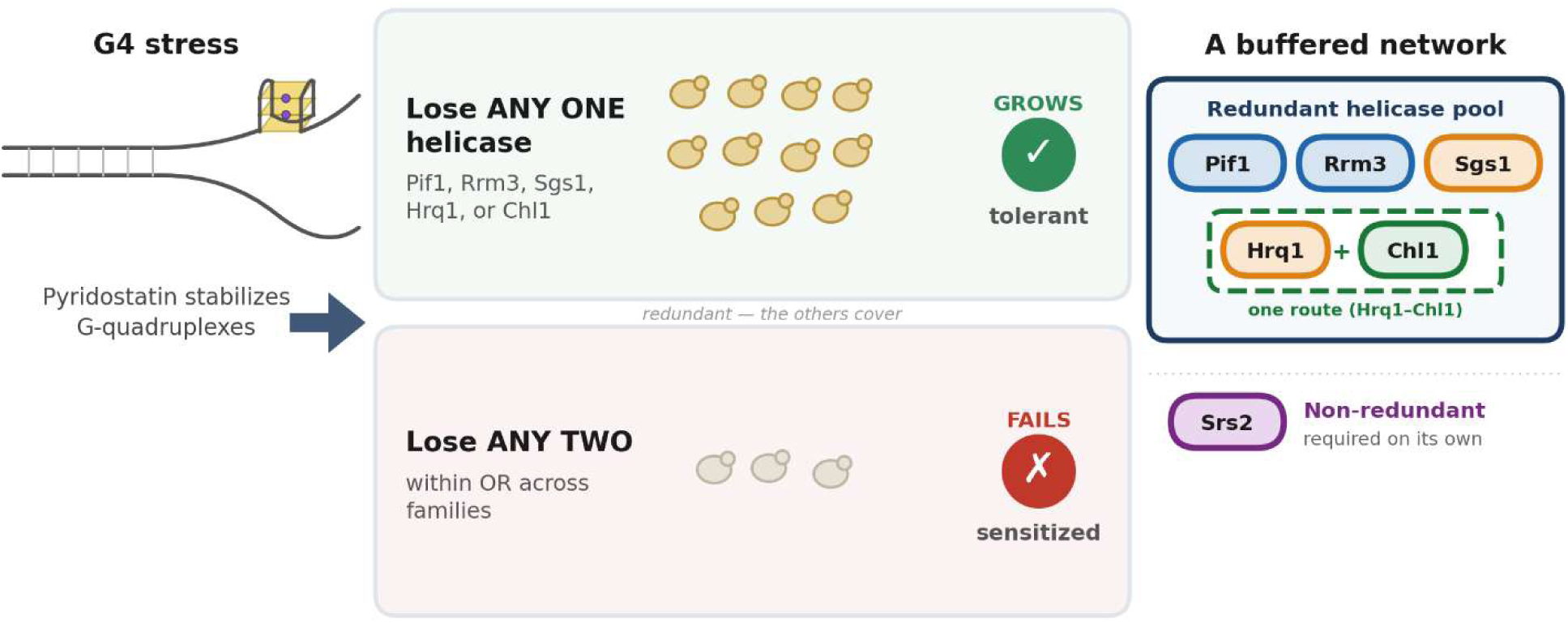

**ARTICLE SUMMARY:** DNA helicases, enzymes that unwind DNA, are thought to dismantle G-quadruplexes (G4s), four-stranded DNA structures that can block DNA metabolism and destabilize genomes. In *Saccharomyces cerevisiae*, we used the chemical pyridostatin to stabilize G4s and measured the growth of helicase mutants. Losing any single helicase had no effect, but losing two together – even from different helicase families – impaired growth. The protein Chl1 works with the helicase Hrq1 in one shared pathway, while Srs2 is uniquely required on its own. G4 tolerance therefore depends on a redundant network of helicases. These findings interest researchers studying genome stability and related human cancer-predisposition disorders.

## INTRODUCTION

Guanine-rich single-stranded DNA (ssDNA) can fold into four-stranded G-quadruplex (G4) structures stabilized by stacked guanine tetrads (Bochman et al. 2012). Putative G4-forming sequences are non-randomly distributed across eukaryotic genomes – enriched at telomeres, replication origins, promoters, and other regulatory loci – and G4 formation has been linked to both normal regulatory functions and to replication fork stalling and genome instability (Rhodes and Lipps 2015; Hansel-Hertsch et al. 2017). Because unresolved G4s threaten genome integrity, cells deploy DNA helicases to unwind or otherwise process them (Lopes et al. 2011; Mendoza et al. 2016; Varshney et al. 2020).

Two helicase families have been the focus of much of this research. The PIF1 family of 5′→3′ helicases, Pif1 and its paralog Rrm3 in the budding yeast *Saccharomyces cerevisiae* (Bochman et al. 2010), promotes replication through G4 motifs and suppresses G4-associated instability (Paeschke et al. 2013; Paeschke et al. 2011; Ivessa et al. 2003).

The RecQ family of 3′→5′ helicases is represented by Hrq1 and Sgs1 in in yeast (Lu and Davis 2021). Sgs1 is an ortholog of the human BLM helicase, and they are both reported to unwind G4 substrates *in vitro* and limit G4-linked recombination and instability *in vivo* (Sun et al. 1998; Huber et al. 2002; Chatterjee et al. 2014; van Wietmarschen et al. 2018). *S. cerevisiae* Hrq1 is the ortholog (Barea et al. 2008) of and functional homolog (Bochman et al. 2014; Nickens et al. 2018; Rogers et al. 2017; Rogers et al. 2020a; Rogers and Bochman 2017) of human RECQL4, mutations in which cause the cancer-predisposition disorders Rothmund-Thomson, RAPADILINO, and Baller-Gerold syndromes (Kitao et al. 1999). Despite extensive *in vitro* characterization (Rogers et al. 2017; Papageorgiou et al. 2023; Li et al. 2023), the relative in vivo contributions of these helicases to G4 tolerance, and the extent to which they act redundantly with Sgs1/BLM, remain incompletely defined (Sanders et al. 2020; Simmons et al. 2021).

The small molecule pyridostatin (PDS) binds and stabilizes G4 structures and has been widely used to probe G4 biology, including the induction of G4-dependent DNA damage (Rodriguez et al. 2012; Rodriguez et al. 2008). PDS therefore offers a tractable means to impose acute, G4-specific genomic stress and to read out the genetic requirements for surviving it. We reasoned that comparing the PDS sensitivity of PIF1- and RecQ-family *S. cerevisiae* mutants would reveal which family is more important for G4 maintenance *in vivo*. However, our results led us away from that framing. We found that single PIF1 and RecQ mutants are uniformly PDS tolerant and that PDS sensitivity is a property of mutant combinations – within and across families – implying functional redundancy rather than family-specific dominance. Pursuing whether this redundancy is a general property of G4-linked helicases, we extended the analysis to Chl1 (the DDX11/ChlR1 ortholog, mutated in Warsaw Breakage Syndrome; (van der Lelij et al. 2010)) and to the anti-recombinase Srs2 (Krejci et al. 2003; Veaute et al. 2003), which is functionally equivalent to human RTEL1 (Barber et al. 2008; Frizzell et al. 2014). Both of these *S. cerevisiae* helicases chosen due to the links to G4 metabolism of their human counterparts (Lerner et al. 2020; Bharti et al. 2013; Vannier et al. 2012; Wu et al. 2020). These experiments both generalized the redundancy and exposed its limits, yielding a buffered network view of G4 tolerance that we develop below.

## MATERIALS AND METHODS

### Yeast strains and growth media

All strains are derivatives of *Saccharomyces cerevisiae* YPH499 (*MATa ura3-52 lys2-801_amber ade2-101_ochre trp1Δ63 his3Δ200 leu2Δ1*). The *pif1-m2* allele was used for all *PIF1* perturbations to preserve mitochondrial Pif1 function (Schulz and Zakian 1994); all other mutations are complete open-reading-frame deletions. Genotypes of all single and combinatorial mutants were confirmed by PCR and Sanger sequencing. The genotypes of all strains used are listed in Table S1. Details of strain construction are available upon request. Yeast cells were grown in YPD (1% w/v yeast extract, 2% w/v peptone, and 2% w/v dextrose).

### Genome-wide G4 prediction and topology

Putative G4-forming sequences were identified across the *Saccharomyces cerevisiae* S288C reference genome (NCBI assembly GCF_000146045.2_R64) using a canonical putative-quadruplex-sequence search – four G-tracts of ≥ 3 guanines separated by loops of 1–25 nt, unioned with strict two-tetrad motifs (G4Hunter score ≥ 1.2; (Bedrat et al. 2016)) – yielding 1,044 loci, which were annotated by genomic feature (ORF, subtelomere, promoter, intergenic, ARS, rDNA, tRNA, and mitochondrial) from the R64 annotation. For each locus, folding topology (parallel, antiparallel, or hybrid) was predicted from primary sequence using G4ShapePredictor under its K⁺ model (Liew et al. 2024).

### AlphaFold3 structure prediction

Each putative G4 structure predicted above was modeled with a local installation of AlphaFold3 (v3.0.3) (Abramson et al. 2024), run via Docker image on a single NVIDIA RTX 4090 GPU. Inputs were bare DNA plus *n* = (number of G-tetrads − 1) channel K+ ions. Every locus was folded in two conditions, apo and with pyridostatin supplied as an additional ligand, from its SMILES string, producing 1,044 matched apo/+PDS pairs (2,088 models). Each job used one model seed with five diffusion samples, with the top-ranked model per condition retained. For each model, we recorded the predicted Template Modeling (pTM) score from the summary confidence output and the mean per-atom predicted Local Distance Difference Test (pLDDT) score over the DNA atoms as a per-locus model confidence. We also computed the apo→+PDS change (ΔpTM and ΔG4-pLDDT) for each matched pair. The generated G4 structures are available in Supplementary File S1.

### PDS sensitivity assays

PDS (APExBIO) was dissolved in DMSO to 10 mM, and aliquots were stored at -20°C. The half-maximal inhibitory concentration (IC_50_) in wild-type cells was determined across a 0.25–32 µM concentration series (Fig. S1). For mutant-panel experiments, cells were grown in the presence of 5 µM PDS (approximating the wild-type IC_50_) or an equivalent volume of DMSO as a solvent control. Growth was monitored in a BioTek Synergy H1 plate reader by monitoring optical density at 600 nm (OD_600_) over 48 h at 30°C with shaking. For each strain, nine biological replicates were assayed in each condition, with DMSO and PDS wells paired by replicate.

### Growth quantification

We quantified cell growth as described in (Sausen and Bochman 2021). Briefly, for each well, growth was summarized as the mean OD_600_ over 48 h to avoid stochastic growth curve effects. Relative growth was computed per biological replicate as the paired ratio of growth in PDS to growth in DMSO (PDS/DMSO), a normalization that removes constitutive differences in baseline fitness among strains so that PDS sensitivity is not confounded by slow growth in the absence of drug. The IC_50_ was determined by fitting dose-response curves (n = 3) via nonlinear least-squares regression using a four-parameter logistic model with the upper and lower asymptotes constrained to 100% and 0%, respectively (Fig. S1). Nonlinear regression was performed in Python 3.12 using the curve_fit function in SciPy (Virtanen et al. 2020), and the figure was generated using Matplotlib. Growth was normalized to the ‘No PDS’ control.

### Statistical analyses

Normality (Shapiro–Wilk; (Shapiro and Wilk 1965)) and homogeneity of variance (Levene; (Brown and Forsythe 1974)) were assessed per group. Because PDS-condition variances were strongly unequal across strains, between-strain differences in relative growth were evaluated within each experimental batch by the Welch-type Alexander–Govern test (Alexander and Govern 1994) with Games–Howell post-hoc comparisons (Games and Howell 1976) against the batch wild-type. Baseline (DMSO) growth was compared across strains analogously. Within-strain DMSO *vs*. PDS differences were tested by paired *t*-tests with Holm correction (Holm 1979). Effect sizes are reported as Hedges’ *g* (Hedges 1981). Wild-type controls differed between experimental batches; each mutant was therefore compared only to the wild-type assayed in the same batch, and family-wise error was controlled within batch. Statistical analyses were performed in Python (SciPy; (Virtanen et al. 2020)); full per-strain results are provided in Table S2. Significance is denoted as * *P* < 0.05, ** *P* < 0.01, *** *P* < 0.001, and **** *P* < 0.0001.

## RESULTS

### PDS raises AlphaFold3 confidence for G4s genome wide

Before treating cells with PDS, we sought to establish a structural footing for the treatment itself. Predicted G4s are distributed throughout the yeast genome (Capra et al. 2010), and PDS is a well-characterized G4-binding ligand (Rodriguez et al. 2012; Rodriguez et al. 2008). We assembled an atlas of 1,044 putative G4-forming loci across the *S. cerevisiae* S288C (R64) genome – 636 within open reading frames, 194 in subtelomeric regions, 137 in promoters, and the remainder in intergenic, ARS, mitochondrial, rDNA, and tRNA regions – and folded each locus with AlphaFold3 (Abramson et al. 2024) as a matched pair in the absence and presence of PDS supplied as a ligand (Fig. 1; Table S3). Including PDS in the input raised AlphaFold3’s confidence in the folded G4 structures genome-wide: the predicted TM-score increased at 96% of loci (ΔpTM = +0.29 ± 0.22; mean 0.25 → 0.54), and the mean per-residue confidence across the G4 increased at 88% (ΔG4-pLDDT = +10.3 ± 9.4; mean 62 → 72). The gain was uniform and present across all three sequence-predicted topologies (parallel, hybrid, antiparallel; ΔG4-pLDDT +10 to +12) (Fig. 1A-C), as well as every genomic feature class (+6 to +19) (Fig. 1D-E), and only weakly related to G4 architecture (|r| ≤ 0.15 with G4Hunter score (Bedrat et al. 2016), tetrad count, or apo confidence) (Table S3). pTM and G4-pLDDT are AlphaFold3 confidence scores (Abramson et al. 2024), so appending a bound ligand raises model confidence without, by itself, constituting a per-locus measurement of thermodynamic stabilization. The direction of increased pTM and pLDDT is exactly the direction that existing PDS biology predicts: it is a well-established G4-stabilizing ligand (Rodriguez et al. 2012; Rodriguez et al. 2008), and a genome-wide increase in structural confidence when PDS is present is the expected *in silico* signature of that activity. Our AlphaFold3 modeling shows PDS consistently engaging G4s of every class across the yeast genome, and it yields a matched apo/+PDS structural set spanning that landscape (per-locus responses and structures in Table S3 and the accompanying data package). The *in vivo* consequences of this predicted G4 stabilization are the subject of the remainder of this study.

**Figure 1.**
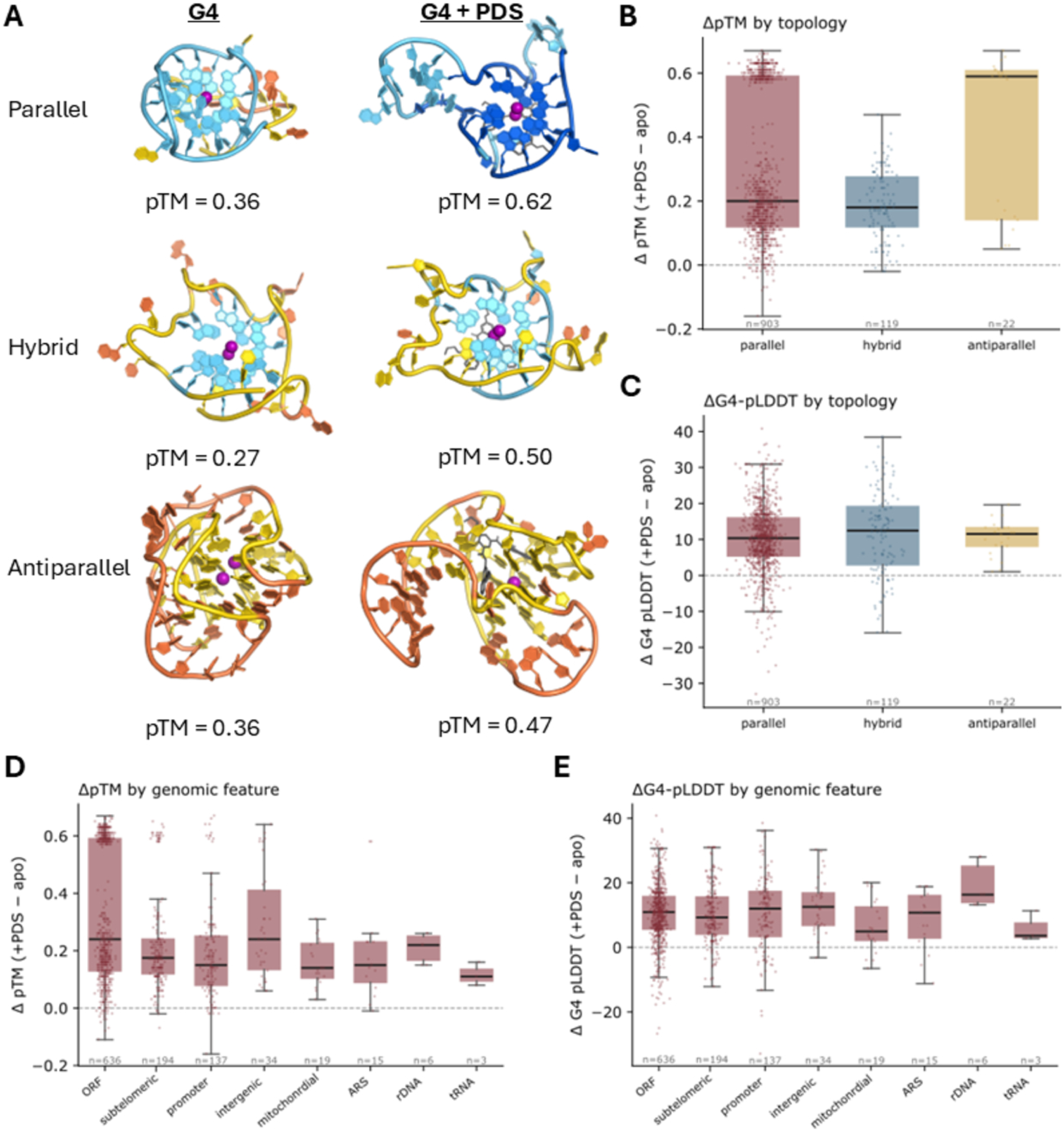
PDS is predicted to stabilize G4 structures genome wide. (A) AlphaFold3 models of putative *S. cerevisiae* G4s with and without PDS bound (Abramson et al. 2024). AlphaFold3 models are colored by per-residue pLDDT (B-factor). K⁺ ions are purple spheres, and PDS is gray. The addition of PDS raises both pTM (shown) and pLDDT. The three rows indicate sequence topology determined using G4ShapePredictor (Liew et al. 2024). Top row: yg4_promXIII (promoter, chr XIII), 5’-GGGAGATAAGAGGGAGCAGGGTGGGGT-3’. Middle: yg4_telI (telomere, chr I), 5’-TGGGTGTGGTGTGGGTGTGGGTGTGGGT-3’. Bottom: atl_0290 (promoter, chr V), 5’-GGGGAGAGGAGGGCTGCGGTTAAGGCTTATGGGGTGTATGGG-3’. PDS raises AlphaFold3 structural confidence in terms of both ΔpTM (B) and ΔG4-pLDDT (C) across G4 topologies (n = 1,044). This confidence response in ΔpTM (D) and ΔG4-pLDDT (E) is uniform across the genomic features listed on the x-axes.

### PDS inhibits wild-type growth with a low-micromolar IC_50_

To establish a principled working dose, we first measured the dose–response of wild-type cells to PDS. Across a concentration gradient covering three orders of magnitude, growth declined sigmoidally with increasing PDS concentration, yielding an IC_50_ of 4.85 µM (95% CI, 4.19–5.50 µM; Fig. S1). We selected 5 µM PDS for all subsequent experiments because dosing at/near the IC_50_ places the assay in the most sensitive region of the dose–response curve, maximizing the power to detect genotype-dependent shifts in either direction.

### Combined loss of PIF1- and RecQ-family helicases sensitizes cells to PDS

We next compared the PDS sensitivity of PIF1- and RecQ-family mutants. Contrary to the expectation that loss of a canonical G4 helicase would confer sensitivity, neither *pif1-m2* nor *rrm3Δ* (Pif1 family), nor *hrq1Δ* or *sgs1Δ* (RecQ family) strains grew significantly worse than wild type in PDS, whether assessed as relative growth or by comparison to wild type (Fig. 2A-D, Table S2). In every case the single-mutant relative growth was statistically indistinguishable from wild type (1.1–1.3× wild-type relative growth; all *P* > 0.15).

**Figure 2.**
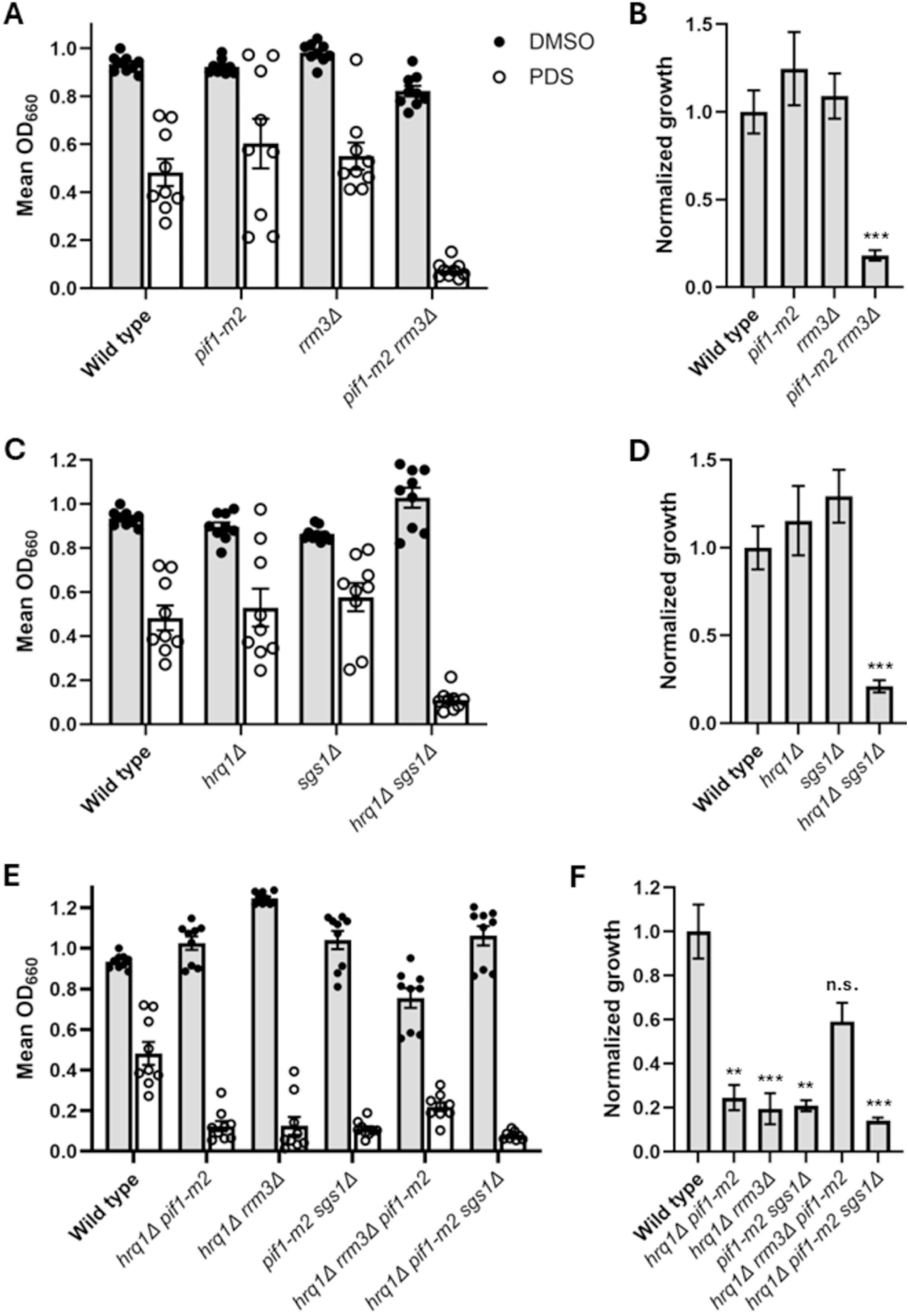
No single PIF1- or RecQ-family helicase is required to maintain fitness during PDS treatment. Growth of wild type and PIF1-family (A), RecQ-family (C), and mixed-family (E) helicase mutant strains in media containing PDS or the solvent control (DMSO). (B, D, F) Relative growth (PDS/DMSO) for wild type and mutant strains. Single mutants are tolerant; the double and triple mutants are sensitized. Independent data points (n = 9) are shown in A, C, and E; bars show the mean ± SEM in all panels. Significance *vs*. wild type was determined by Games–Howell analysis. We defined significance as \*\*\*\**P* < 0.0001, \*\*\**P* < 0.001, \*\**P* < 0.01, and \**P* < 0.05; n.s., not significant.

Sensitivity emerged only when two helicases were lost together. The within-family double mutants *pif1-m2 rrm3Δ* and *hrq1Δ sgs1Δ* were both strongly sensitized to PDS, retaining only 0.18× and 0.21× the wild-type relative growth, respectively (*P* = 5.6 × 10⁻⁴ and *P* = 6.8 × 10⁻⁴; Fig. 2A-D, Table S2). Critically, the requirement was not confined within families: the cross-family double mutants *hrq1Δ pif1-m2* (0.25× wild type; *P* = 0.002), *hrq1Δ rrm3Δ* (0.20×; *P* = 9.0 × 10⁻⁴), and *pif1-m2 sgs1Δ* (0.21×; *P* = 0.002) were each sensitized to a comparable degree (Fig. 2E-F, Table S2). Thus, a cell can lose any one of these four helicases and tolerate G4 stabilization. However, loss of two, regardless of family boundary, falls below the threshold required for normal growth. These data indicate that G4 tolerance is buffered by the combined activity of multiple, partially interchangeable helicases rather than by either family alone.

This buffering extended to higher-order combinations, with one informative exception. The triple mutant *hrq1Δ pif1-m2 sgs1Δ* was among the most strongly sensitized strains in the panel (0.14× wild-type relative growth; *P* = 1.0 × 10⁻³; Fig. 2E-F, Table S2), consistent with the loss of a third buffering activity. In contrast, *hrq1Δ pif1-m2 rrm3Δ* was not significantly sensitized (0.59× wild type; *P* = 0.13; Fig. 2E-F), despite lacking on additional helicase relative to the double mutants from which it derives. We attribute this apparent discordance to our previous findings that *hrq1Δ pif1-m2* and *hrq1Δ pif1-m2 rrm3Δ* cells exhibit a lower rate of gross chromosomal rearrangements (GCRs) than *pif1-m2* and *pif1-m2 rrm3Δ* cells alone, respectively (Nickens and Bochman 2022), indicating that loss of Hrq1 suppresses a genome-instability phenotype caused by PIF1 helicase mutation(s). A triple mutant built on this *hrq1Δ pif1-m2* core inherits that suppression, accounting for its relative tolerance to G4 stabilization. The contrasting behavior of the two triples – one sensitized, one buffered by an internal genetic suppression – reinforces that the relevant unit is the combination of activities present, not simply the number of helicases lost.

### The redundancy extends to Chl1, which is epistatic with Hrq1

The data above suggest that it is not PIF1-family helicases or RecQ-family helicases that are the most important for G4 maintenance *in vivo* but, rather, the combined action of enzymes in both families buffering against PDS stress. Is this a trait that is specific to just these two helicase families though? If PDS tolerance reflects a distributed helicase capacity rather than a property of two specific families, then other G4-linked helicases might participate in the same buffering network. We therefore tested Chl1, the yeast ortholog of human DDX11/ChlR1 (van der Lelij et al. 2010; Lerner et al. 2020). As with the canonical single mutants, the *chl1Δ* strain was not sensitized to PDS (Fig. 3A-B). When combined with Pif1- or RecQ-family mutations, *chl1Δ* reproduced the redundancy pattern: *chl1Δ pif1-m2*, *chl1Δ rrm3Δ*, and *chl1Δ sgs1Δ* each shifted toward reduced relative growth (0.63–0.71× wild type; Hedges *g* = −0.7 to −1.4), and the Chl1-block relative-growth comparison was significant overall (*P* = 0.004; Table S2). However, we note that the individual *chl1Δ* double mutants were variable and did not reach per-strain significance against wild type (*P* = 0.11–0.62). We therefore report these as a consistent directional trend rather than as individually significant interactions (Table S2).

**Figure 3.**
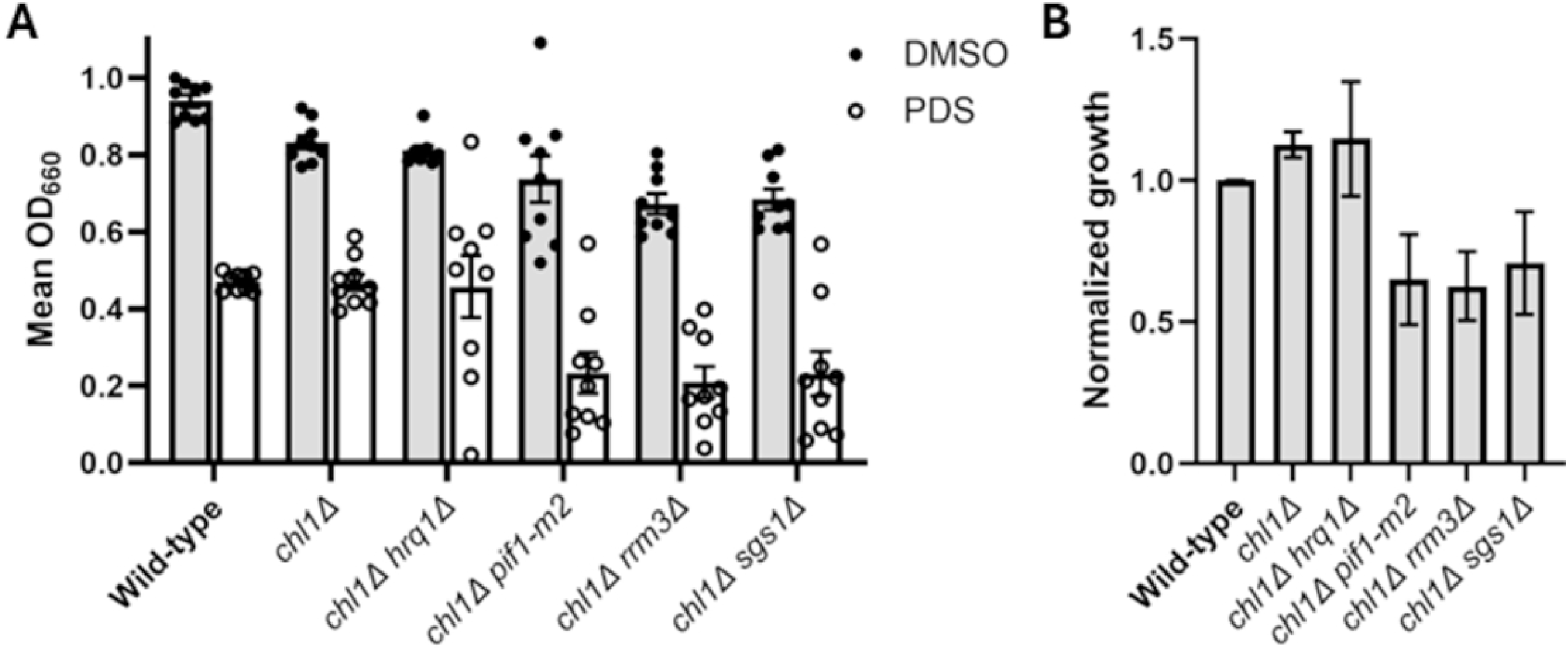
Chl1 mutants display the same pattern. (A) Growth of wild type and strains lacking *CHL1* in media containing PDS or the solvent control (DMSO). (B) Relative growth (PDS/DMSO) for wild type and mutant strains. The data were graphed and analyzed as in Figure 2.

Notably, one combination behaved differently. The *chl1Δ hrq1Δ* strain was not sensitized to PDS; its relative growth matched wild type (1.15× wild type; *P* = 0.97, *g* = +0.33). This is the predicted signature of two genes acting in a single epistasis group: whereas *chl1Δ* is synthetic with each of the other three helicase mutations, it shows no interaction with *hrq1Δ* – the same interaction profile displayed by *hrq1Δ* itself. The well-powered tolerance of *chl1Δ hrq1Δ* (*i.e.*, the absence of a synthetic interaction) suggests that Chl1 and Hrq1 function in a shared route, with the more variable sensitized *chl1Δ* doubles providing consistent directional support that Chl1 engages the broader helicase network.

### Srs2 is uniquely required as a single mutant

The redundancy observed for the helicases above predicts that no single helicase deletion should confer PDS sensitivity. However, Srs2, the functional analog of human RTEL1 (Barber et al. 2008; Frizzell et al. 2014; Vannier et al. 2012; Wu et al. 2020), violated this prediction. The *srs2Δ* strain was the only single mutant in the entire panel that was significantly sensitized to PDS, displaying only 0.25× the wild-type relative growth (*P* = 1.1 × 10⁻⁴; Fig. 4A-B, Table S2). Combining *srs2Δ* with *pif1-m2*, *hrq1Δ*, or *chl1Δ* did not produce growth meaningfully worse than *srs2Δ* alone, consistent with the single mutant already approaching the floor of the assay (Fig. 4A-B, Table S2). Because Srs2 cannot be compensated by any of the helicases tested, it defines a non-redundant requirement that is distinct from the buffered helicase pool – a separate functional axis rather than another interchangeable member of it.

**Figure 4.**
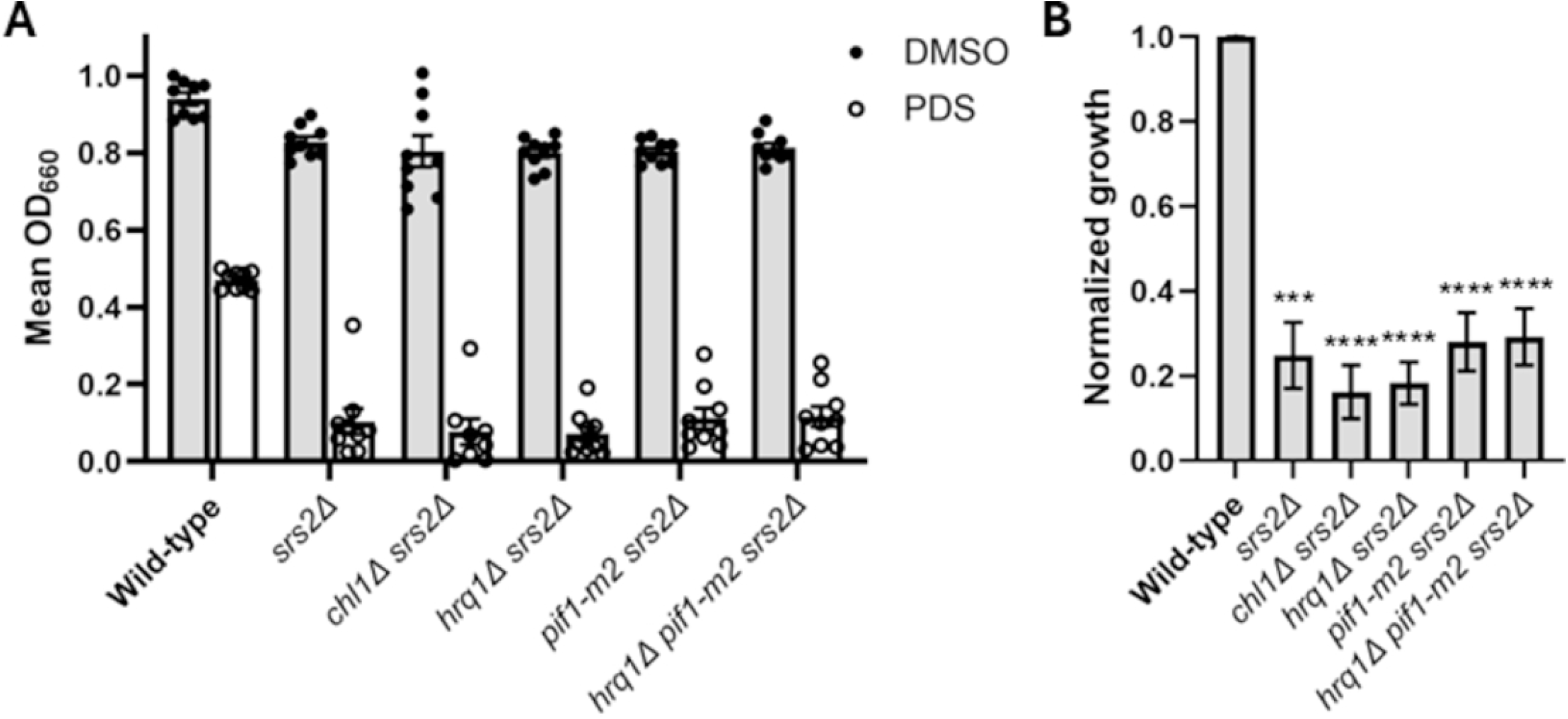
Srs2 is required for PDS resistance. (A) Growth of wild type and strains lacking *SRS2* in media containing PDS or the solvent control (DMSO). (B) Relative growth (PDS/DMSO) for wild type and mutant strains. The data were graphed and analyzed as in Figure 2.

### An integrated genetic interaction map

Synthesizing the panel, we constructed a genetic interaction map in which helicases are nodes and synthetic PDS sensitivity defines edges (Fig. 5). Five interactions are statistically supported (the within- and cross-family Pif1/RecQ doubles); the *chl1Δ* interactions are directionally consistent but individually weak and are drawn as such; *chl1Δ hrq1Δ* is rendered as an epistatic pair with no connecting edge; and Srs2 sits apart as a sensitive single mutant with no redundant partner. The map captures the central architecture: a redundant, cross-family helicase pool – within which Hrq1 and Chl1 constitute one route – together with a non-redundant Srs2 requirement. Notably, there are several helicase mutant combinations that we could not test due to synthetic lethality: *rrm3Δ sgs1Δ*, *rrm3Δ srs2Δ*, and *sgs1Δ srs2Δ* (Schmidt and Kolodner 2004).

**Figure 5.**
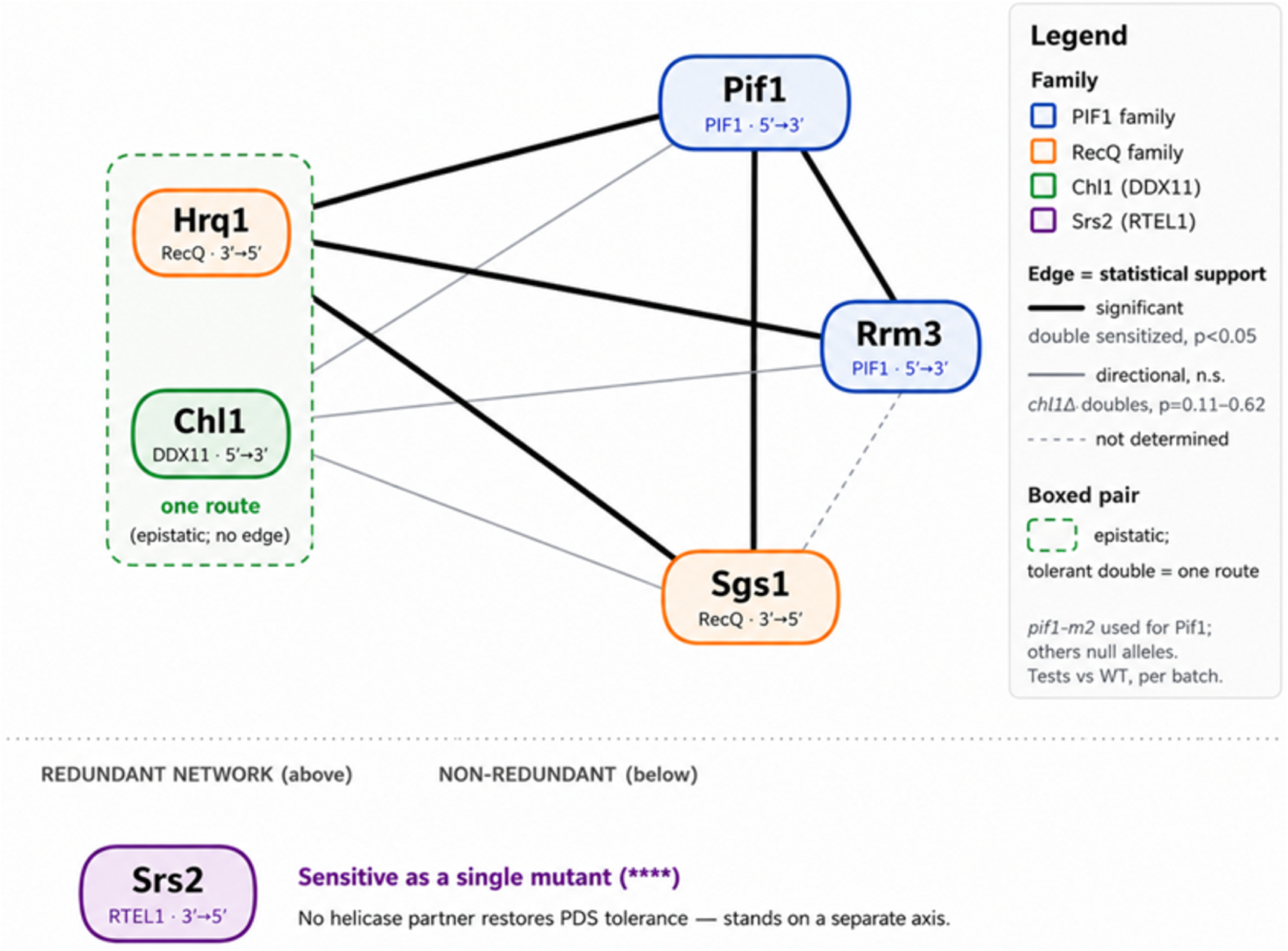
Genetic interaction map of PDS sensitivity. Nodes are helicases; edge thickness encodes statistical support for synthetic sensitivity (thick = significant double-mutant sensitization *vs*. wild type; thin = directional but not significant; dotted = not determined). Hrq1 and Chl1 are boxed as one epistatic route (no connecting edge); Srs2 is shown separately as a non-redundant sensitive single mutant.

## DISCUSSION

We set out to compare the relative importance of PIF1- and RecQ-family helicases for G4 maintenance and instead found that, at the level of growth under G4 stabilization, the historical narrative surrounding the importance of these enzyme families is miscast (*e.g.*, (Paeschke et al. 2013)). No single helicase among Pif1, Rrm3, Hrq1, Sgs1, or Chl1 is required; sensitivity arises only during the combined loss of two of these helicase, and it crosses both family boundaries and DNA translocation polarities. The most parsimonious interpretation is that these helicases contribute to a shared, buffered capacity to resolve or bypass G4-associated genomic stress, such that wild-type cells retain enough total activity to tolerate PDS even when any one enzyme is missing. Because PDS engages predicted G4s broadly – spanning every topology and genomic feature class, from promoters and open reading frames to telomeres and tRNA/rDNA loci (Fig. 1) – this buffered capacity is best read as a genome-wide property rather than the specialization of a few sites.

This conclusion refines rather than refutes the substantial literature implicating PIF1- and RecQ-family helicases in G4 metabolism (Paeschke et al. 2013; Paeschke et al. 2011; Ivessa et al. 2003; Sun et al. 1998; Huber et al. 2002; Chatterjee et al. 2014; van Wietmarschen et al. 2018; Rogers et al. 2017; Papageorgiou et al. 2023; Li et al. 2023). These enzymes remain important for G4 maintenance. Our data simply indicate that their *in vivo* contributions to surviving acute G4 stabilization are mutually redundant, so that the phenotypic requirement is revealed only in combination. Such redundancy is a plausible reason that single-helicase mutants have often shown modest or context-dependent phenotypes in prior work (Paeschke et al. 2011; van Wietmarschen et al. 2018; Lerner et al. 2020; Mendoza et al. 2016; Varshney et al. 2020).

The placement of Chl1 and Hrq1 in one epistasis route is notable because the two enzymes are otherwise dissimilar: Chl1 (DDX11/ChlR1) is a 5′→3′ iron-sulfur helicase tied to replication-coupled sister-chromatid cohesion (Hirota and Lahti 2000; Pisani et al. 2018; Petronczki et al. 2004; Skibbens 2004), whereas Hrq1 (RECQL4) is a 3′→5′ RecQ-family enzyme (Rogers et al. 2020b). Their cooperation in a single functional route – rather than acting as independent, interchangeable members of the pool – raises the possibility of sequential or coupled action on a common substrate, a hypothesis that warrants direct biochemical testing (for example, whether one stimulates the other’s activity on G4 substrates). Because both human orthologs are genome-instability and cancer-predisposition genes, a conserved Chl1–Hrq1 functional relationship would be of broad interest. Indeed, both enzymes are already known to function in DNA inter-strand crosslink repair in *S. cerevisiae*, though in different pathways (Rogers et al. 2017). Perhaps G4 structures represent another shared *in vivo* substrate.

Srs2 stands apart from this redundant logic. As the only single mutant sensitized to PDS, Srs2 performs a function that no helicase in the pool can replace. Srs2 is best known not as a G4 resolvase but as an anti-recombinase that strips Rad51 from single-stranded DNA and dismantles toxic recombination intermediates (Krejci et al. 2003; Veaute et al. 2003). We therefore favor a two-layer model (Fig. 6): a redundant helicase pool resolves or bypasses PDS-stabilized G4s at sites where they form in single-stranded DNA (e.g., replication forks), while a non-redundant Srs2 function manages the recombinogenic consequences of the DNA breaks that nonetheless arise. A direct prediction follows: if *srs2Δ* PDS sensitivity reflects toxic Rad51-dependent recombination, then deleting *RAD51* should suppress it. Testing this hypothesis via *rad51Δ* epistasis analyses, together with assessing the effects of catalytic-dead alleles to dissect which Hrq1 activity supports the Chl1 route, are a priority moving forward.

**Figure 6.**
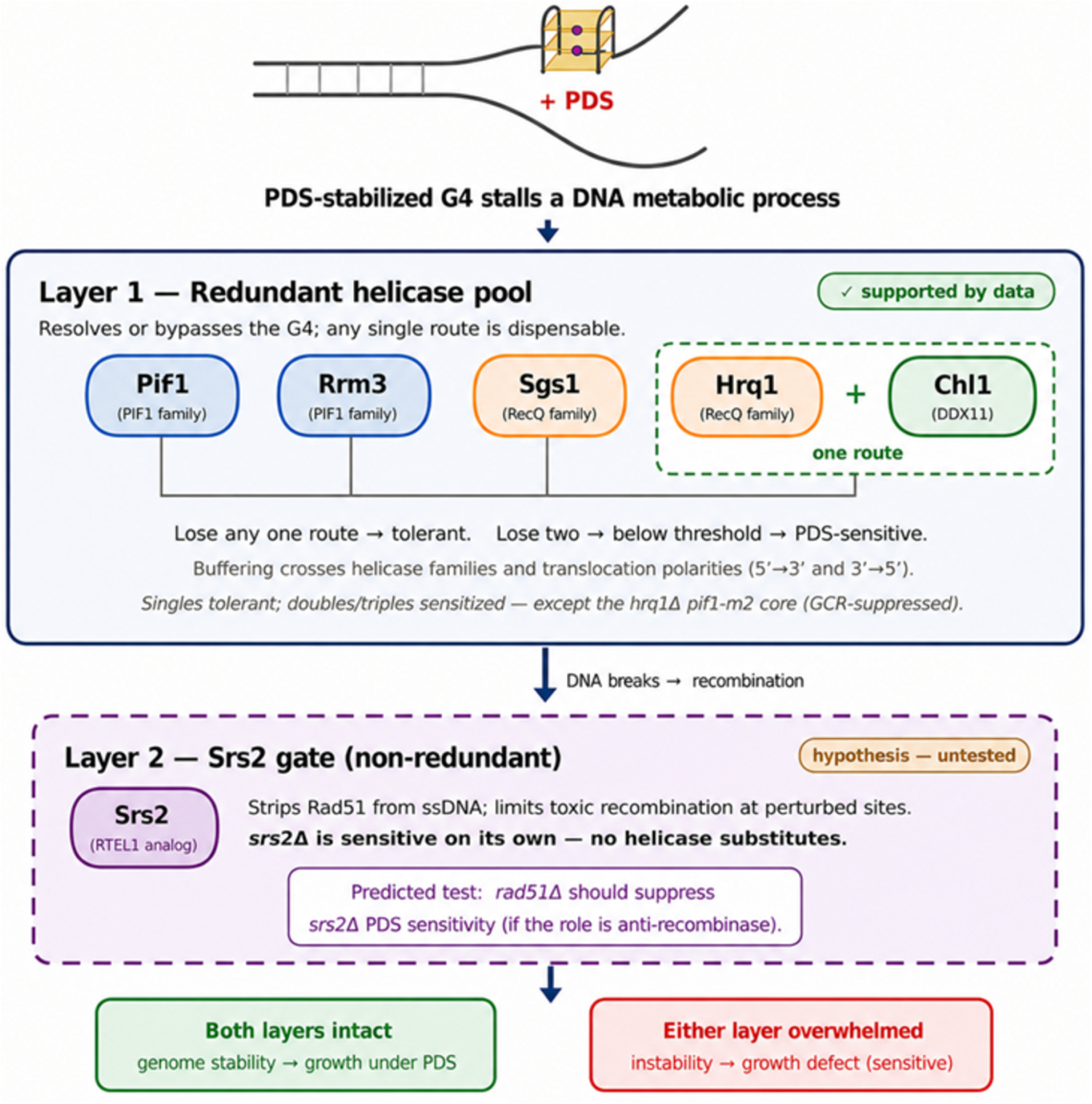
Two-layer model of helicase buffering to survive G4 stabilization. Our data indicate a redundant helicase pool (Pif1, Rrm3, Sgs1, and the Hrq1–Chl1 route) resolves or bypasses PDS-stabilized G4s at sites where the structures form in ssDNA (*e.g.*, DNA replication forks). A non-redundant Srs2 function – hypothetically, its role in recombination – manages the DNA repair consequences of DNA breaks at stabilized G4s. A predicted *rad51Δ*-suppression test is indicated.

Several caveats bound our conclusions. First, our readout is growth under chronic chemical G4 stabilization; tolerance at the level of fitness does not exclude sub-lethal genome instability in single mutants, and our ‘no effect’ statements apply to the growth endpoint. Second, we have extended the redundancy beyond the original four helicases to Chl1 and Srs2, which is sufficient to show that the pattern is not idiosyncratic to the PIF1 and/or RecQ helicase families but does not establish that redundancy is a universal property of helicases. Third, PDS is considered a pan-G4 stabilizing ligand, but to our knowledge this has not been proven conclusively within *S. cerevisiae*. Our AlphaFold3 atlas (Fig. 1 and File S1) provides an *in silico* signature consistent with genome-wide engagement: including PDS raised model confidence at the great majority of loci and across every topology and feature class. Nonetheless, pTM and pLDDT are confidence scores, not measurements of thermodynamic stabilization or *in vivo* occupancy (Abramson et al. 2024), so this remains a prediction rather than direct proof. Confirming engagement biophysically, and testing an orthogonal G4 ligand (*e.g.*, PhenDC3 (De Cian et al. 2007)) with our mutant strains, would further strengthen these analyses. Within these bounds, however, the data support a coherent reframing: G4 maintenance in budding yeast is a distributed, buffered network of redundant helicase activities, atop which a non-redundant Srs2 function guards against the recombinogenic fallout of G4-perturbed genome maintenance.

## DATA AVAILABILITY

Strains are available upon request. The authors affirm that all data necessary for confirming the conclusions are present within the article, figures, and supplemental material. Supplemental materials include Figure S1, Tables S1-S3, and File S1. Supplemental material is available online.

## ACKNOWLEDGMENTS

We thank members of the Bochman Lab for insightful discussions and feedback pertaining to this work, especially Michael Kumcu and his statistical queries. We are also grateful to Google LLC for providing local access to AlphaFold3.

## STUDY FUNDING

This work was funded by the College of Arts and Sciences, Indiana University.

## CONFLICT OF INTEREST

The authors declare that they have no conflicts of interest.

